# Glacier ice archives fifteen-thousand-year-old viruses

**DOI:** 10.1101/2020.01.03.894675

**Authors:** Zhi-Ping Zhong, Natalie E. Solonenko, Yueh-Fen Li, Maria C. Gazitúa, Simon Roux, Mary E. Davis, James L. Van Etten, Ellen Mosley-Thompson, Virginia I. Rich, Matthew B. Sullivan, Lonnie G. Thompson

## Abstract

While glacier ice cores provide climate information over tens to hundreds of thousands of years, study of microbes is challenged by ultra-low-biomass conditions, and virtually nothing is known about co-occurring viruses. Here we establish ultra-clean microbial and viral sampling procedures and apply them to two ice cores from the Guliya ice cap (northwestern Tibetan Plateau, China) to study these archived communities. This method reduced intentionally contaminating bacterial, viral, and free DNA to background levels in artificial-ice-core control experiments, and was then applied to two authentic ice cores to profile their microbes and viruses. The microbes differed significantly across the two ice cores, presumably representing the very different climate conditions at the time of deposition that is similar to findings in other cores. Separately, viral particle enrichment and ultra-low-input quantitative viral metagenomic sequencing from ∼520 and ∼15,000 years old ice revealed 33 viral populations (i.e., species-level designations) that represented four known genera and likely 28 novel viral genera (assessed by gene-sharing networks). *In silico* host predictions linked 18 of the 33 viral populations to co-occurring abundant bacteria, including *Methylobacterium*, *Sphingomonas*, and *Janthinobacterium*, indicating that viruses infected several abundant microbial groups. Depth-specific viral communities were observed, presumably reflecting differences in the environmental conditions among the ice samples at the time of deposition. Together, these experiments establish a clean procedure for studying microbial and viral communities in low-biomass glacier ice and provide baseline information for glacier viruses, some of which appear to be associated with the dominant microbes in these ecosystems.

**Importance:** This study establishes ultra-clean microbial and viral sampling procedures for glacier ice, which complements prior *in silico* decontamination methods and expands, for the first time, the clean procedures to viruses. Application of these methods to glacier ice confirmed prior common microbiological findings for a new ice core climate record, and provides a first window into viral genomes and their ecology from glacier ice across two time horizons, and emphasizes their likely impact on abundant microbial groups. Together these efforts provide clean sampling approaches and foundational datasets that should enable simultaneous access to an archived virosphere in glacier ice.

## Introduction

The first reports of microbes in glacier ice appeared early in the 20^th^ century (1, 2), but were largely ignored until microorganisms were investigated in the deep Vostok ice core in the 1980s (3). This motivated further studies of glacier ice microbes near the end of the last century [reviewed in (4–7)]. These reports indicated that microbial biomass is very low in most glacier ice samples, with the estimated number of microbial cells ranging from 10^2^ to 10^4^ cells ml^−1^ (5) [compared to, for example, 10^4^–10^6^ cells ml^−1^ in marine water (8)]. The microbes found in glacier ice cores are typically interpreted to represent the microbes in the atmosphere at the time of their deposition, and hence reflect climatic and environmental conditions during that time period (4, 9). Taxonomically, *Proteobacteria*, *Actinobacteria*, *Firmicutes*, and *Bacteriodetes* are the dominant bacteria found in ice cores (5, 10–12), including some that have been successfully cultured from long frozen glacier ice (13–17). Most of these isolates are psychrotolerants (17, 18), which have optimal growth temperatures well above freezing, but can be preserved under cold environments such as glacier ice for a long time (19). Although there is currently no direct evidence for *in situ* activity, several studies have hinted at the possibility of microbial activity in frozen glacier ice based on the detection of some excess gases (e.g., CO2, CH4, and N2O), which may be produced by post-depositional microbial metabolisms (20–22).

Viruses are the most numerous constituents of microbial communities in oceans, with an abundance of 10^6^ to 10^9^ particles ml^−1^ of seawater (23, 24). They can alter microbial communities through lysis, horizontal gene transfer, and metabolic reprogramming (25–31). Typically, some abundant viruses detected in Arctic sea ice and ancient cryopegs were previously predicted to infect the dominant microbial members in the community and to likely modulate host adaptations to extreme cold and salt conditions (Z. P. Zhong, Z. Rapp, J. M. Wainaina et al., submitted for publication). If the non-polar marine and polar cryosphere systems are any indication, then viruses archived in glacier ice are also likely to have infected the co-occurring microbial hosts, whether before and/or after ice formation. Most ice core microbiological studies have focused on microbial communities and how to use them to understand past climatic and environmental conditions archived in the glaciers (4–7). In contrast, there are only two reports of viruses in glacier ice. One detected the atmosphere-originated tomato-mosaic-tobamovirus RNA in a 140,000-year-old Greenland ice core by reverse-transcription polymerase chain reaction amplification (32), and the other reported the presence of virus-like particles (VLPs) deep (i.e., 2749 and 3556 meters depth) in the Vostok ice core using transmission electron microscopy (4). Notably, viruses preserved in glacier ice have yet to be studied using a modern viral ecogenomics toolkit (29), nor have any viral genomes or long viral contigs been reported.

Although microbes in glacier ice have been recognized since early in the 20^th^ century and investigated more rigorously after the 1980s, it remains challenging to use sequencing approaches to study microbial or viral communities in such low-biomass and remote environments. This is primarily due to the low quantity of nucleic acids that can be extracted. In addition, because of this low biomass, contaminations from sampling, storage, and processing are a major issue for studying microbial communities in glacier ice (33). Considerable efforts have been made to develop clean sampling techniques, and several surface decontamination strategies have been published that describe the removal of potential contaminants on ice core surfaces before melting them for microbial analysis (34, 35). These methods efficiently eliminate suspected contaminants on the core surfaces and have been widely adopted for microbial investigations of glacier ice [e.g., (18, 22, 36, 37)]. However, the efficiency of these decontamination methods varies with the decontamination instruments, laboratory personnel, and details of the procedures. Therefore, we sought to establish clean procedures to remove microbial contaminants on ice surfaces by creating and experimenting on several sterile artificial ice core sections covered with known bacteria, viruses, and free DNA (see Materials and Methods). Then we applied these new procedures, together with our previously described *in silico* decontamination methods for removing suspected contaminants introduced during the processing of ice in the laboratory (12), to investigate microbial and viral communities archived in two ice cores drilled on the summit and plateau of the Guliya ice cap in northwestern China (35°17’ N; 81°29’ E; ∼6650 m asl).

## Results and Discussions

### Establishing clean surface-decontamination procedures with mock contaminants

In the field, no special procedures are used to avoid microbial contamination during ice core drilling, handling, and transport. Therefore, ice core surfaces contain microbial contaminants that impede the identification of microbial communities archived in the ice (35, 38). To develop a clean surface-decontamination procedure for removing possible microbial contaminants on the ice core surfaces and for collecting clean ice for microbial investigations, we constructed sterile artificial ice core sections, and covered them with a known bacteria (*Cellulophaga baltica* strain 18, CBA 18), virus (*Pseudoalteromonas* phage PSA-HP1), and free DNA (from lambda phage) [see Materials and Methods & Fig. 1; and as performed in (35)]: The decontamination procedure involved three sequential steps, including scraping (i.e., cutting) with band saw and then washings with clean ethanol and water, to remove the outermost ice layers (a total of ∼1.5 cm of the core radius; see Materials and Methods and Fig. 1, and Fig. S1 in the supplemental material at https://doi.org/10.6084/m9.figshare.11427246.v1). The ice removed during each step and the remaining inner ice were collected to determine if the contaminants (including bacteria, viruses, and free DNA) were removed completely (see Materials and Methods & Fig. 1).

**Figure 1.**
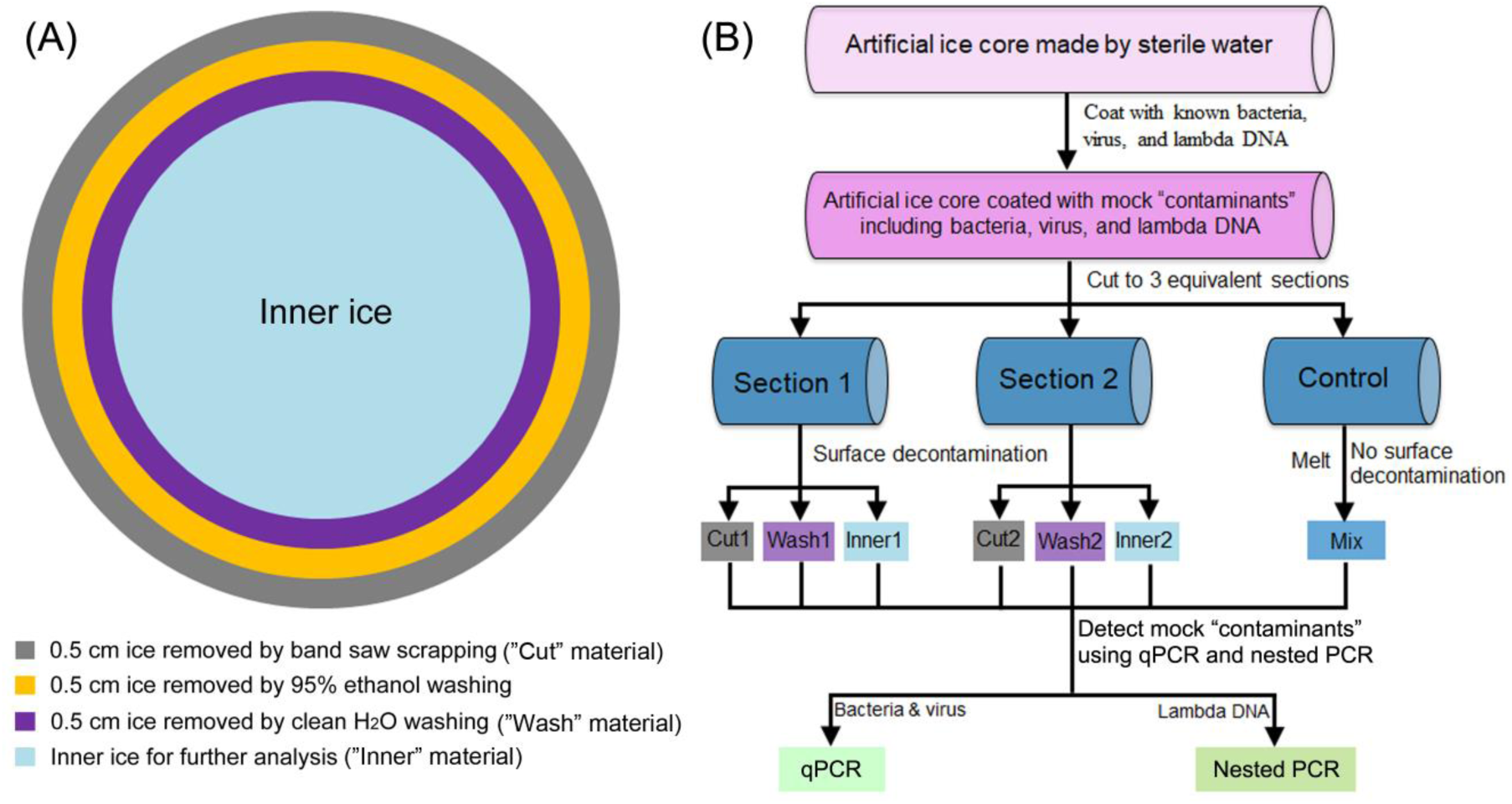
Overview of decontamination protocol. (**A**) Schematic of layered removal of the outer core surface to obtain clean inner ice. (**B**) Experimental approach establishing decontamination procedures using sterile artificial ice core sections coated with mock “contaminants”. Cut, Wash, and Inner represent ice samples collected from band saw scrapping, water washing, and the inner ice, respectively. Mix represents a sample from the melted ice of a control ice core section prepared without decontamination processing. The mock contaminants were detected by qPCR and nested PCR (see Methods).

The bacterial and viral contamination in each sample was quantified using strain-specific primers and qPCR (see Materials and Methods). The results showed orders of magnitude reduction in contaminant bacteria and viruses after being processed with the surface decontamination procedure described above (Fig. 2A). Nested PCR was used to detect the contaminant lambda phage DNA, which was absent in the inner ice (Fig. 2B). These results indicate that the decontamination procedure removed contaminants such as bacteria, viruses, and free DNA from the surface ice and left clean ice for further microbial and viral analysis. Clean ice was also successfully obtained for microbial investigation as reported in a previous methods paper (35). However, we constructed different decontamination systems (Fig. S1) and expanded the clean procedures to also decontaminate viral particles from glacier ice core surfaces.

**Figure 2.**
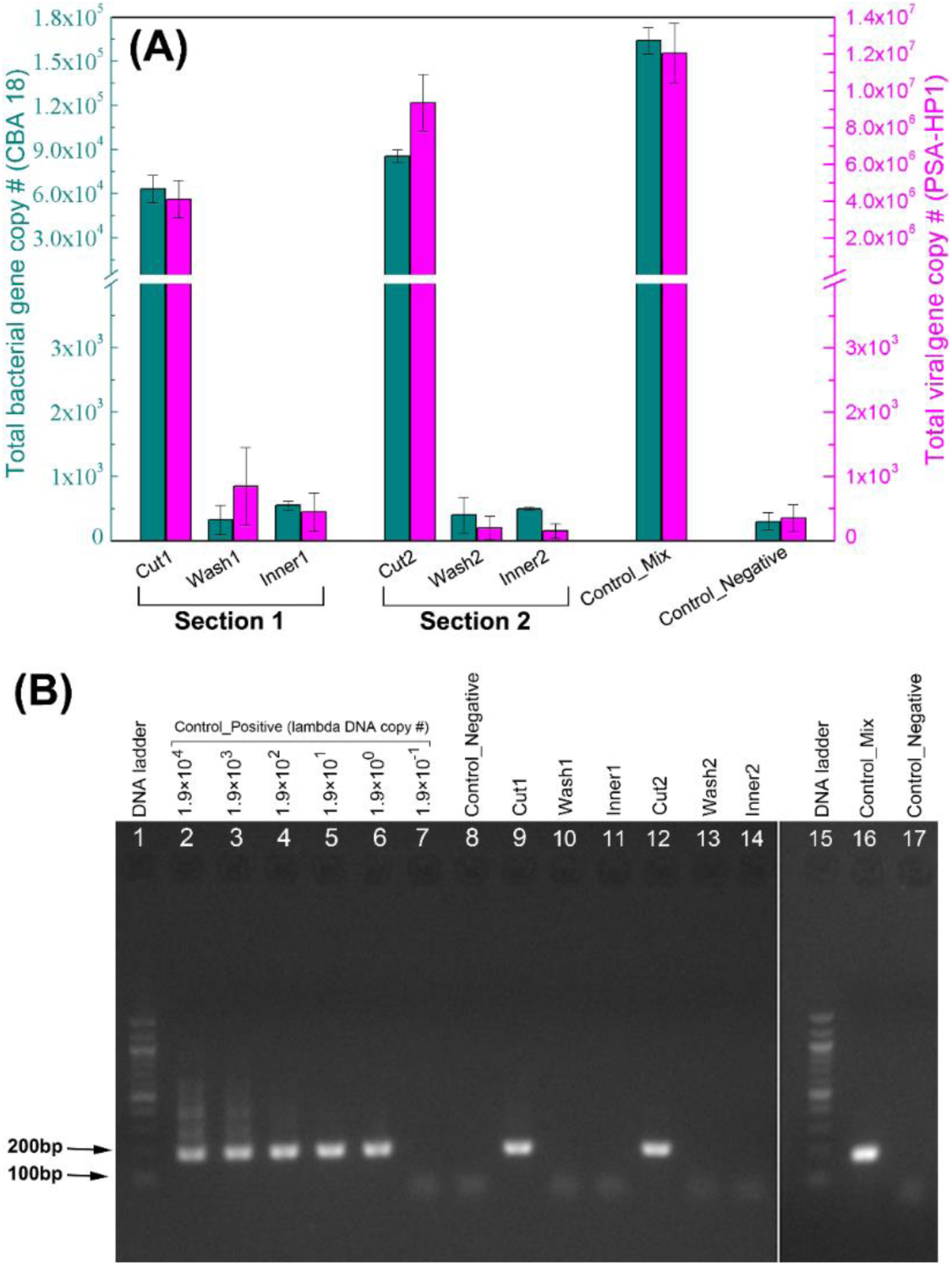
Examination of the effectiveness of decontamination procedures. (**A**) Total bacterial (dark teal color) and viral (purple color) numbers were quantified by qPCR using strain-designed primers in all samples collected in Fig. 1. (**B**) Lambda DNA was detected using nested PCR with designed outer and inner primer sets for lambda DNA. PCR products from inner primer sets were visualized by agarose gel electrophoresis; 1, 100bp DNA ladder; 2–7 represent 1.9×10^4^, 10^3^, 10^2^, 10^1^, 10^0^, and 10^−1^ (10-times dilution from standards) copies of lambda DNA, respectively, used as templates for nested PCR; 8, Control_Negative (no template); 9, Sample Cut1; 10, Wash1;11, Inner1; 12, Cut2; 13, Wash2; 14, Inner2; 15, 100bp DNA ladder (same as 1); 16, Control_Mix; 17, Control_Negative (same as 8). Names of all ice samples are coded as described in Fig. 1.

### Decontamination method provides clean ice from glacier core sections

After we established that the surface-decontamination procedure removed surface contaminants, we then used authentic ice core sections to further evaluate the procedure. Two sections (Samples PS.D13.3 and PS.D13.5 from 13.34–13.50 and 13.50–13.67 m depth, respectively), obtained from a shallow ice core (PS ice core) drilled in 1992 from the plateau of the Guliya ice cap (Fig. 3), were decontaminated using the procedures described above (Fig. 1). The ice removed during each step (Cut: saw-scraped ice; Wash: H2O-washed ice), and the inner ice (Inner) for each section was collected as described above (see Materials and Methods & Fig. 1). Microbial profiles of six samples (three samples — Cut, Wash, and Inner — from each of the two ice sections) were investigated using Illumina Miseq 16S rRNA gene amplicon sequencing. The quality-controlled data were rarefied to 12,000 sequences in each sample (i.e., each MiSeq sequencing library) for further analysis.

**Figure 3.**
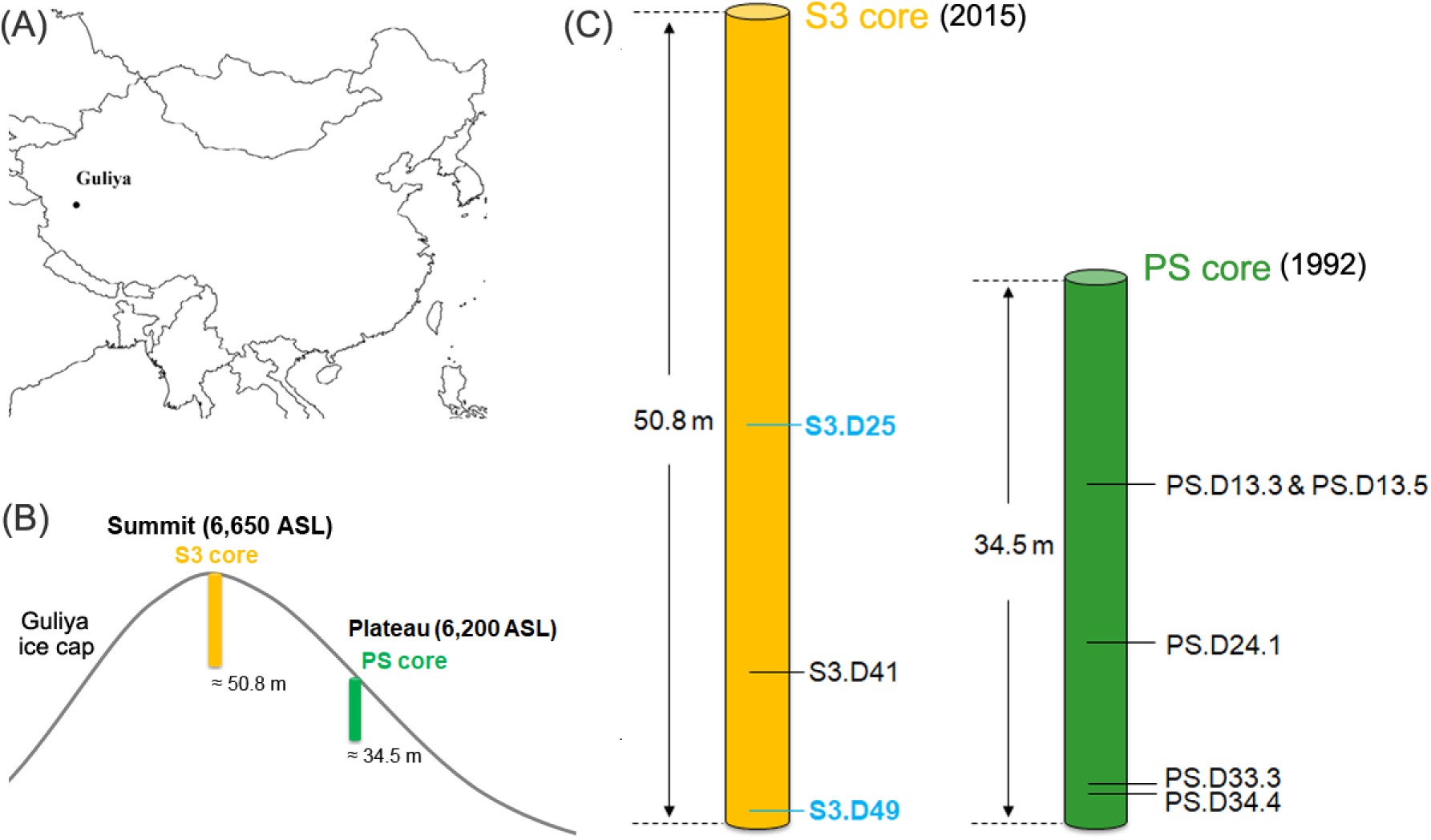
Location of the Guliya ice cap (**A**), drilling sites of the S3 and PS ice cores (**B**), and sampling depths of eight ice samples for investigating the microbial and viral communities (**C**). S3 and PS cores were drilled from the summit and plateau of Guliya ice cap, respectively (**B**). The drill data and length of two ice cores are indicated (**C**). The sample names are coded by core and depth, e.g., for PS.D13.3, PS is the ice core name, and D13.3 indicates a depth of 13.3 m below the surface. All samples were subjected to microbial investigations, and two samples S3.D25 and S3.D49 (light blue) were selected for viral investigation.

The 14 most abundant classes of microbes, each of which accounted for ≥0.5% of the sequences in at least one sample, comprised 98.9% of the total 72,000 sequences in the six samples. These groups were designed as “major classes” and were selected to compare the microbial communities of all Cut, Wash, and Inner samples for both ice sections (Fig. S2A). Within each ice section, the most abundant classes were shared across the Cut, Wash, and Inner samples (Fig. S2A). For example, all the top seven of the most abundant classes *Actinobacteria*, *Cytophagia*, *Flavobacteriia*, *Sphingobacteriia*, *Alphaproteobacteria*, *Betaproteobacteria*, and *Deltaproteobacteria* were represented in the three (i.e., Cut, Wash, and Inner) PS.D13.3 samples; these classes comprised 92.7%, 96.5%, and 97.9% of the microbial communities in the Cut, Wash, and Inner samples, respectively (Fig. S2A). These results suggest that the surface ice was only slightly contaminated and that the contaminants were not abundant and diverse enough to alter the overall microbial community composition based on the most abundant microbial groups in these ice core sections. The PS ice core was drilled in 1992 using an electromechanical drill with no drilling fluid (39); in general the surfaces of these cores are less contaminated than ice cores extracted using a fluid in the borehole (38). Some microbial classes that were detected in the Cut and Wash samples with low abundance (e.g., *Bacilli* and *Gammaproteobacteria*) were nearly absent in the Inner sample (Fig. S2A), suggesting these were surface contaminants that were successfully removed by the decontamination procedure. Similarly, we compared the microbial communities at the genus level for all of the samples from the same two ice core sections (Fig. S2B). The Cut, Wash, and Inner samples in the same ice core section shared their most abundant genera (Fig. S2B) while some minor genera were removed after surface decontamination and considered as contaminants (e.g., *Bacillus* and *Vibrio*; Fig. S2B). This finding is consistent with previous reports that some *Bacillus* strains were contaminants in glacier ice samples [e.g., (12, 38)].

### Microbial profiles differ between the PS and S3 ice cores

Once a clean decontamination procedure was established with authentic ice core sections, we investigated the microbial communities from five different depths (i.e., 13.3, 13.5, 24.1, 33.3, and 34.4 meters) in the PS ice core, and compared them with the communities from the three summit 3 (S3) ice core samples (i.e., S3.D25, S3.D41, and S3.D49) (Fig. 3). These three S3 samples were processed at the same time, and the 16S rRNA gene data for two (i.e., S3.D41 and S3.D49) of them were published previously (12). Four background controls were co-processed with the glacier ice samples to trace the background microbial profiles, which were then proportionally removed *in silico* from amplicon data of the PS ice core samples (see Materials and Methods), according to our previously published method (12).

After the *in silico* decontamination, we compared the microbial community composition between and within ice cores. Reads were rarefied to 24,000 sequences in each sample, and collectively the samples contained 254 bacterial genera, 118 of which were taxonomically identified to the genus level (see Table S1 in the supplemental material at https://doi.org/10.6084/m9.figshare.11427246.v1). The 32 most abundant genera, defined as those comprising at least 0.5% of sequences in at least one ice sample, collectively represented >96.0% of each community (Fig. 4). Genera including *Janthinobacterium* (relative abundance 1.0–23.8%), *Polaromonas* (2.6–4.1%), *Flavobacterium* (2.3–23.6%), and unknown genera within the families *Comamonadaceae* (15.5–24.3%) and *Microbacteriaceae* (7.1–48.5%) were abundant and present in all five PS samples (Fig. 4). This indicates that members belonging to these lineages are adapted to cold environments and may subsist over long periods of time, although their relative abundances vary across ice core depths (ages). These genera and families have also been reported as abundant groups in glacier ice cores in many previous studies [e.g., (5, 10, 12, 40–44)]. The detection of bacterial sequences belonging to similar genera in ice core samples from different glaciers located around the world can be explained by the ubiquitous distribution of certain species in geographically distant environments (45, 46). The three S3 ice core samples shared some abundant genera with the five PS ice core samples, such as *Janthinobacterium*, *Herminiimonas*, and *Flavobacterium* (Fig. 4). However, several abundant genera in the S3 ice core samples were nearly absent in the PS ice core samples, including *Sphingomonas*, *Methylobacterium*, and an unclassified genus in the family *Methylobacteriaceae* (Fig. 4). Thus, there are fundamental differences in the microbial communities between the ice cores retrieved from the plateau (shallow part) and the summit of the Guliya ice cap.

**Figure 4.**
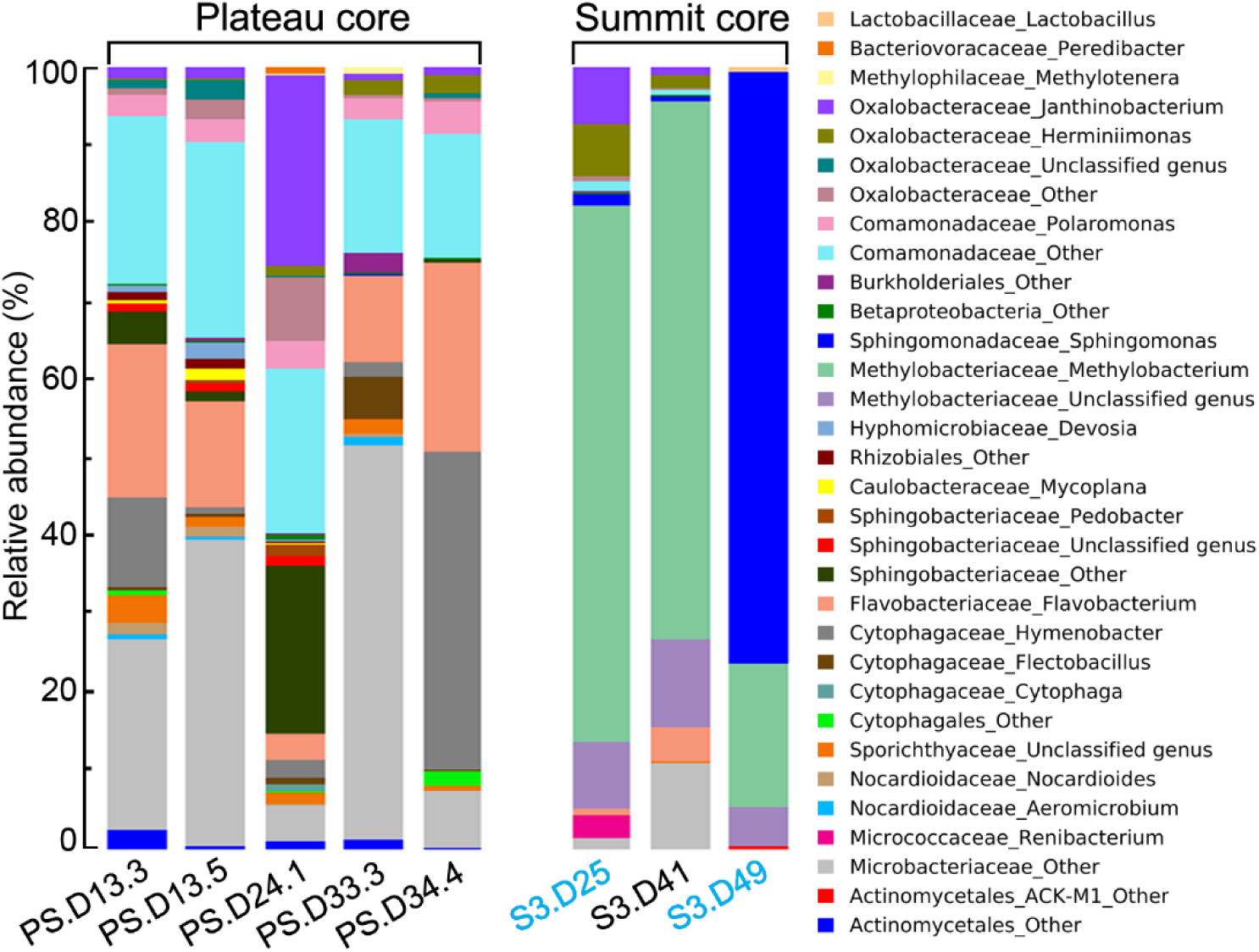
Microbial profiles of the 32 most abundant genera in the PS and S3 ice core samples, as a percent of the total 16S rRNA gene amplicon sequences. The key indicates genera, preceded by family, or order in cases where family is not assigned. Genera labeled “Other” represent sequences with unknown genus-level taxonomy, i.e., distinct from taxonomically-assigned genera in the reference database. The 32 most abundant genera, defined as those comprising at least 0.5% of the sequences in at least one ice sample, collectively represented >96.0% of each community. The total relative abundance of these genera was normalized to 100% in this figure.

We next used principal coordinates analysis (PCoA) to compare microbial community compositions among all eight samples and found that the communities were clustered by the ice core (Fig. 5), separating along the first principle coordinate (which accounted for 68.2% of community variability; the second axis accounted for 13.4%). Analysis of similarity statistics (ANOSIM) confirmed that the microbial communities of samples from the plateau core were significantly different than those from the summit core (p = 0.02, n = 999). The two ice cores were drilled at different elevations (6,200 and 6,650 m) for the plateau and summit cores, respectively; Fig. 3 & Table S2), which represent very different environmental conditions in terms of UV radiation, air temperature, and oxygen concentration (L. G. Thompson, E. Mosley-Thompson et al., unpublished data). These factors undoubtedly influence the microbial communities at the time of deposition. In addition, all PS ice core samples were from the shallower part of the ice cap (top 34.5 m of the ∼310 m thick ice field) (39) and were much younger than the three samples from the S3 core (∼70–300 years versus ∼520–15,000 years old; Table S2), which were collected near the bottom of summit ice core (∼51 m length; Fig. 3).

**Figure 5.**
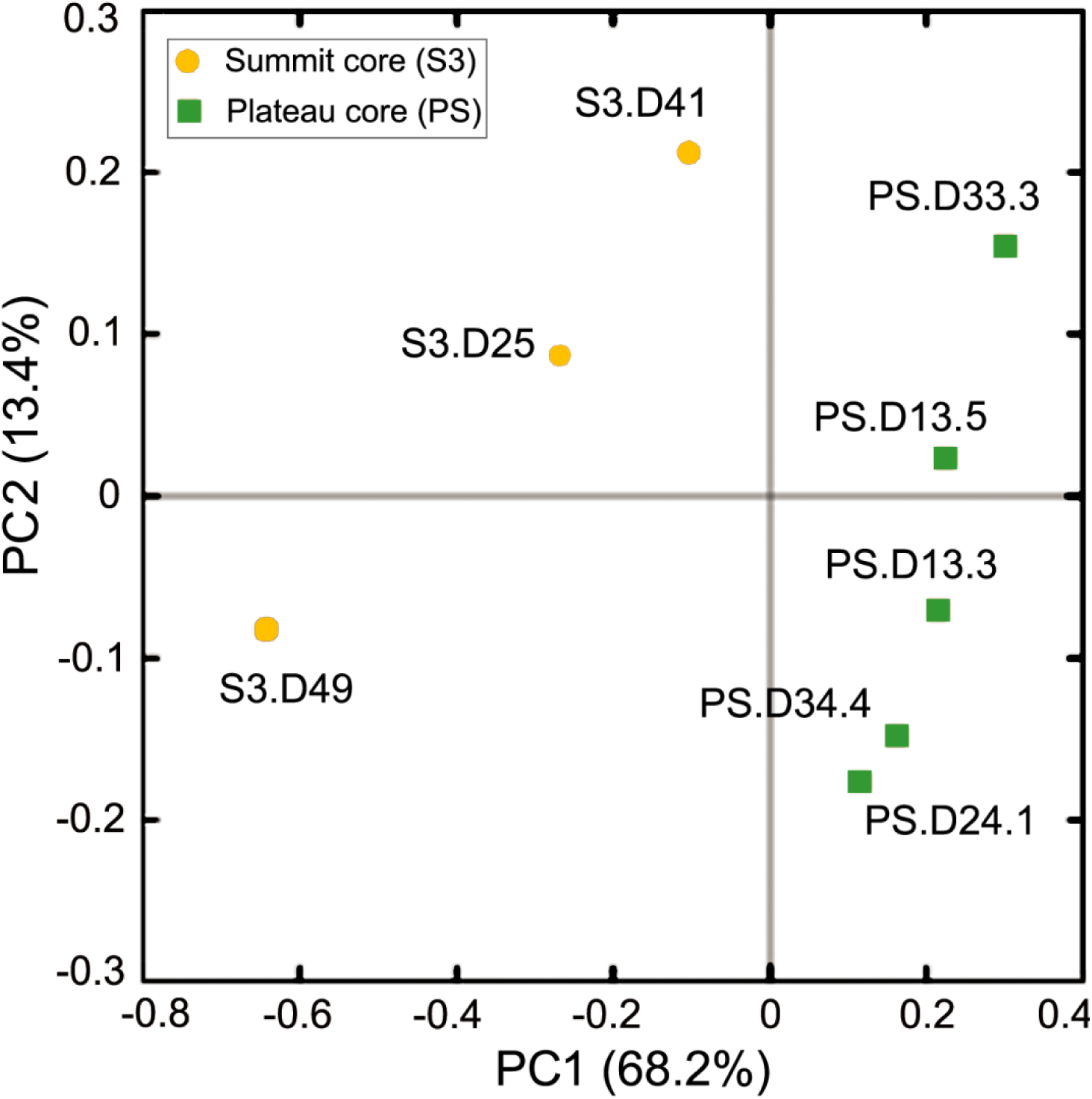
Communities from the two cores (denoted by color) were distinct; sample names are indicated next to each symbol. PCoA was performed on the weighted UniFrac metric, which accounts for the relative abundance and inferred relatedness of the lineages present.

Therefore, the ice samples from the two different ice cores represent very different climate conditions at the time of deposition. This is further illustrated by differences in several environmental parameters (e.g., concentration of insoluble dust and ions such as sulfate and sodium) between the two ice cores (Table S2). To further identify the environmental parameters potentially influencing these microbial communities, two-tailed Mantel tests were performed to examine the relationships between environmental properties (Table S2) and microbial community compositions. Parameters, including elevation, ice age, and concentrations of dust, chloride, sulfate, and sodium significantly (p ≤ 0.05) correlated with microbial community compositions (Table S3). This further supports the above discussion that explains the differences between the microbial communities in the two ice cores, and is consistent with many previous reports that the microbial communities archived in glacier ice often reflect the differences in many physicochemical parameters such as dust concentration (47–49) and some ion concentrations (37, 50). These findings provide empirical support that the ice core microbial communities reflect climate conditions at the time they were deposited.

### Bacteriophage archived in an ice core

Although it is well accepted that diverse microbes are archived in glacier ice (4–7), information about viruses in these habitats is still scarce, mainly due to the low biomass of viruses in glacier ice and the lack of a single and universally shared gene for viruses that can be used for amplicon-based approaches. Advances in sample processing and library generation approaches now enable to study viruses in ultra-low-biomass environments through viral metagenomics (28, 51). We adopted these technologies to investigate viruses in two ice samples (S3.D25 and S3.D49) selected from the S3 ice core. The samples were selected based on their difference in ice age (∼520 versus ∼15,000 years old), climate conditions (colder versus warmer based on the δ^18^O data, not shown), and on dust concentrations, which are up to 10 times higher in the S3.D49 sample (Table S2).

Counts of VLPs in the two samples were below the detection limit using a wet-mount method [<10^6^ VLPs ml^−1^; (52)], so viral metagenomic DNA in these two samples was subjected to low-input quantitative viral metagenomic sequencing as described previously (53–55). After sequencing, quality control, and *de novo* assembly, we obtained 1,849 contigs with a length of ≥10 kb (Table S4). Overall, VirSorter predicted 43 “confident” viral contigs [≥10 kb in size and Categories 1, 2, 4, or 5; Table S4 (56)]. These viral contigs were dereplicated into 33 viral populations using currently accepted cut-offs that approximate species-level taxonomy (28, 57–59). On average, 1.4% (2.2 and 0.6% for S3.D25 and S3.49, respectively) of the quality-controlled reads were recruited to these viral populations (Table S4). Low percentage of reads recruited to predicted viral sequences is not unusual for low-input viromes, and consistent with previous studies including from more diverse communities [e.g., as low as 0.98%, (28, 60)]. While previous studies detected tomato-mosaic-tobamovirus RNA and VLPs in glacier ice (4, 32), this is the first report of viral genome fragments assembled *de novo* from such ecosystem. Rarefaction curves were constructed (see Materials and Methods) and showed that both viromes approached saturation of long viral populations (≥10 kb) at the sequencing depth used in this study (Fig. S3), though we note that low-input libraries have to be PCR-amplified prior to sequencing (15 PCR cycles in this study) and this may overestimate the redundancy within a library due to PCR duplicates.

### Viral communities consist of mostly novel genera and differ between depths

With 33 viral populations (length ≥10 kb) obtained from the two S3 ice samples, we then evaluated how viruses in this unexplored extreme environment compared to known viruses. Because viruses lack a single, universally shared gene, taxonomies of new viruses are now commonly established using gene-sharing analysis from viral sequences longer than 10 kb in length (60, 61). In our dataset, that meant comparing shared gene sets from 33 viral populations with genomes from 2,304 known viruses in the NCBI RefSeq database (version 85; Table S5) using vConTACT version 2 (60, 61). Gene sharing analyses in vConTACT produce viral clusters (VCs), which represent approximately genus-level taxonomic assignments (31, 60–63). Of the 33 viral populations, four were clustered into four VCs with the RefSeq viral genomes, two formed a VC with only ice populations, and the other 27 populations remained isolated as singleton or outlier populations (Fig. 6A; Table S5). Therefore, only four populations (12%) could be assigned a formal taxonomy: they belonged to four different genera in the families *Siphoviridae* (three genera) and *Myoviridae* (one genus) within the order *Caudovirales* (Table S5). These taxonomic results indicated glacier ice supports a diversity of unique viruses, consistent with studies in oceans (52% unique genera) (31) and soils (61% unique genera) (64).

**Figure 6.**
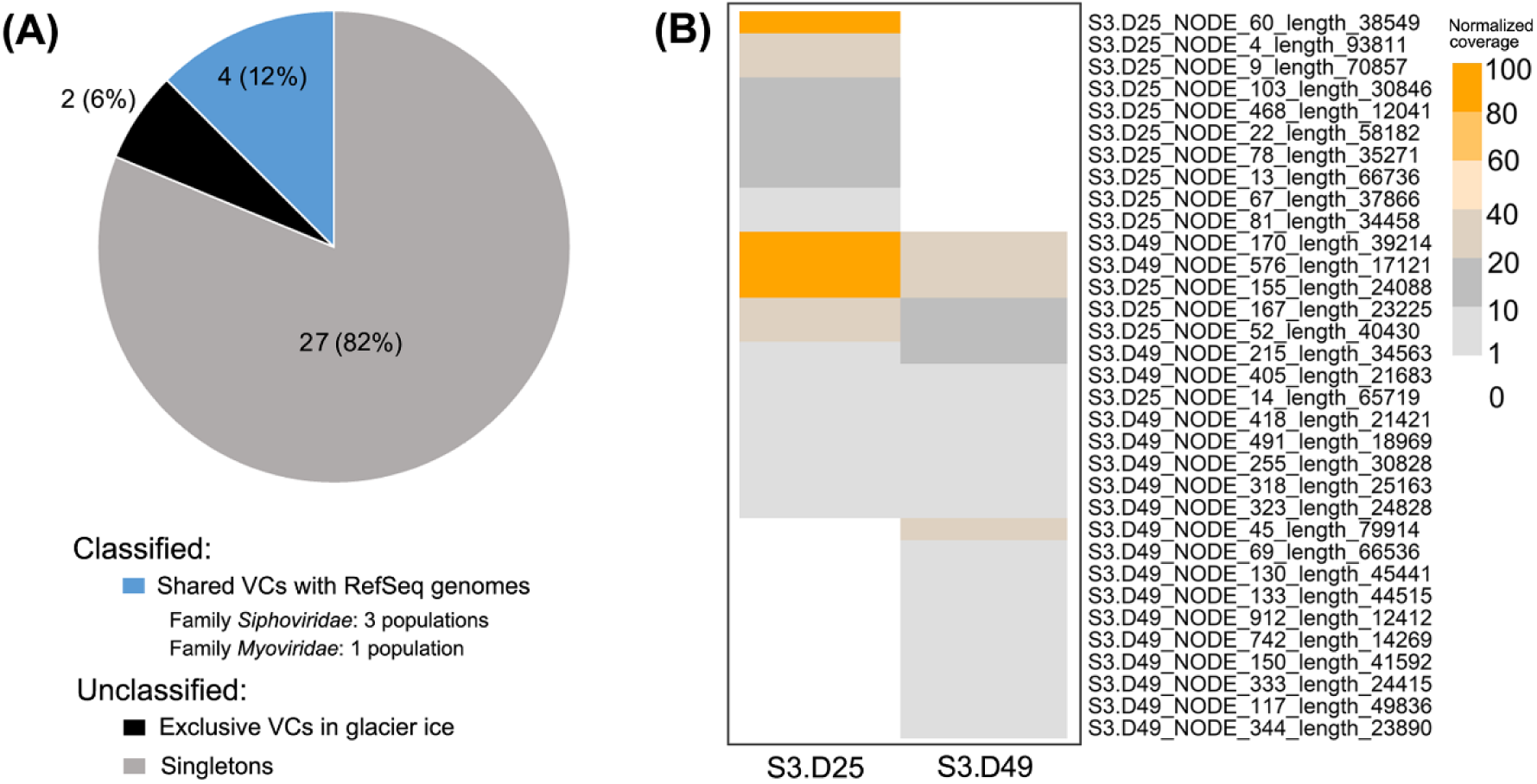
Thirty-three viral populations (≥10 kb) were recovered from two glacier ice samples. (**A**) Viral populations were classified into three groups: ‘Singletons’ (gray) that had no close relatives; ‘Exclusive VCs’ (black) that were viral clusters (VCs) of exclusively glacier ice populations; and ‘Shared VCs’ (blue) which included glacier ice populations and viral genomes from RefSeq. (**B**) The normalized coverage of these 33 viral populations were generated by mapping quality-controlled reads to viral populations, and were normalized to per gigabase of metagenome.

We then looked more closely at the population (∼species) level to compare viral communities archived in two different depths of the S3 ice core. To do this, we first calculated the abundance of each viral population (total = 33) by mapping quality-controlled reads from each sample to the population contigs and then normalizing by gigabase of virome (Table S6; see Materials and Methods). All populations were then used to compare the viral community compositions between the two samples by a heatmap, which showed that the glacier ice from the two depths contained a mix of shared and depth-unique viral populations (Fig. 6B). This was similar to the difference in microbial communities of these samples in which some microbial groups were shared while some varied between the two depths (Fig. 4; Table S1). Previous studies have also reported different microbial community structures in ice samples collected from different depths of the same ice core, which probably reflects differences in the environmental conditions at the time the different ice samples were deposited (48, 65). Interestingly, three viral populations were abundant (relative abundance >10%) in both depths: S3.D49_NODE_170_length_39214, S3.D49_NODE_576_length_17121, and S3.D25_NODE_155_length_24088 (contig names, Fig. 6B; Table S6). This suggests that these viruses may be active in these ice cores, or that a large number of virus particles were initially deposited so that a sufficient amount was still intact for DNA extraction and sequencing after being frozen for fifteen thousand years.

### Viruses predicted to infect dominant microbes in glacier ice

Microbial analysis found that both the S3.D25 and S3.D49 samples were dominated by the bacterial genus *Methylobacterium*, an unclassified genus within the family *Methylobacteriaceae*, and genus *Sphingomonas*, with relative abundances of 18.2–67.5%, 5.0–8.3%, and 1.4–75.3%, respectively (Fig. 4). In addition, the genera *Janthinobacterium* (7.1%) and *Herminiimonas* (6.6%) were also abundant in S3.D25, but were absent or rare (<0.01%) in S3.D49 (Fig. 4). All of these genera are common abundant microbial groups in glacier ice (5, 10, 12, 40–44). In addition, many members belonging to these genera are psychrophilic bacteria and have been revived and isolated from glacier ice (13, 40, 66–68). These results indicate that glacier ice serves as an archive for abundant cold-adapted microbes that might have been active long ago or may still be currently active with the potential to revive and be present in ecosystems after the glaciers melt in the future. Therefore, we next explored the potential impacts of viruses on these abundant microbes by linking viruses to their hosts *in silico*.

Hosts for the 33 viral populations were predicted using three *in silico* methods: similarities in viral and bacterial nucleotide sequences (31, 69), composition (70), or CRISPR spacer matches (31). The sequence similarity method (Blastn) predicted hosts for 14 of the 33 populations (Table S7), whereas the sequence composition method (VirHostMatcher) linked nine populations to microbial hosts (Table S8; see Materials and Methods). Six populations were successfully linked to their hosts by both methods, and these host linkages were of high confidence since the predicted hosts taxonomy was consistent across methods for all six populations (Table S7 & 8). The CRISPR method matched hosts for three viral populations (Table S9), two of which were also linked to hosts by the sequence similarity method but none of them was matched by the sequence composition method (Table S7, 8, & 9). One of the CRISPR-based prediction linked one population (S3.D49_NODE_130_length_45441) to a species within the *Brucella* genus. However, members of this genus was not identified in this sample (i.e., S3.D49; Fig. 4 & Table S1). The same viral population was predicted to infect a *Sphingomonas* host, one of the most abundant members of the microbial community in this sample, through the Blastn method. Hence, this CRISPR-based linkage to *Brucella* was interpreted as a false-positive and discarded for the rest of the analyses. Although only about half (18 of 33 populations) of the viral populations were linked to a host by at least one of the three methods, these host predictions indicated that viruses in glacier ice were infectious at some time (whether before and/or after ice formation) in these extreme cold and high elevation environments, and that they probably played an important role in modulating microbial communities.

The predicted host genera that were most abundant in the same ice cores included *Methylobacterium*, *Sphingomonas*, and *Janthinobacterium* (Fig. 4; Table S1). Many members of these genera are psychrophilic bacteria as mentioned above. The relative abundance of *Methylobacterium*-associated viral populations was high in both S3.D25 (67.5%) and S3.D49 (18.2%; Fig. 7), which was consistent with the dominance (48.2% and 44.0%, respectively) of this bacterial genus in the microbial communities of these two samples (Fig. 7). Similarly, *Janthinobacterium*-linked viruses had a high relative abundance of 7.1% in the S3.D25 sample, whose microbial community was dominated by the genus *Janthinobacterium* with 4.5% relative abundance (Fig. 7); *Sphingomonas*-associated viruses represented 3.1% of communities in the S3.D49 sample, while members of *Sphingomonas* accounted for 75.3% of the microbial profiles in this sample (Fig. 7). The relatively high abundance of these genera and their associated viruses suggests that the viruses we recovered infected abundant microbial groups and thus might play a major role in this extreme ecosystem by influencing their hosts when they are active, although it is still uncertain when the infections occurred. Notably, no host could be predicted for about half of the viral populations, partly due to the limitations of available reference databases and techniques used for host prediction (69). As methods improve and host databases expand [e.g., presently there are 94,759 bacterial genomes in the Genome Taxonomy Database website (71)], continued studies will likely provide more complete understanding of the relationship between viruses and their microbial hosts in the ice cores.

**Figure 7.**
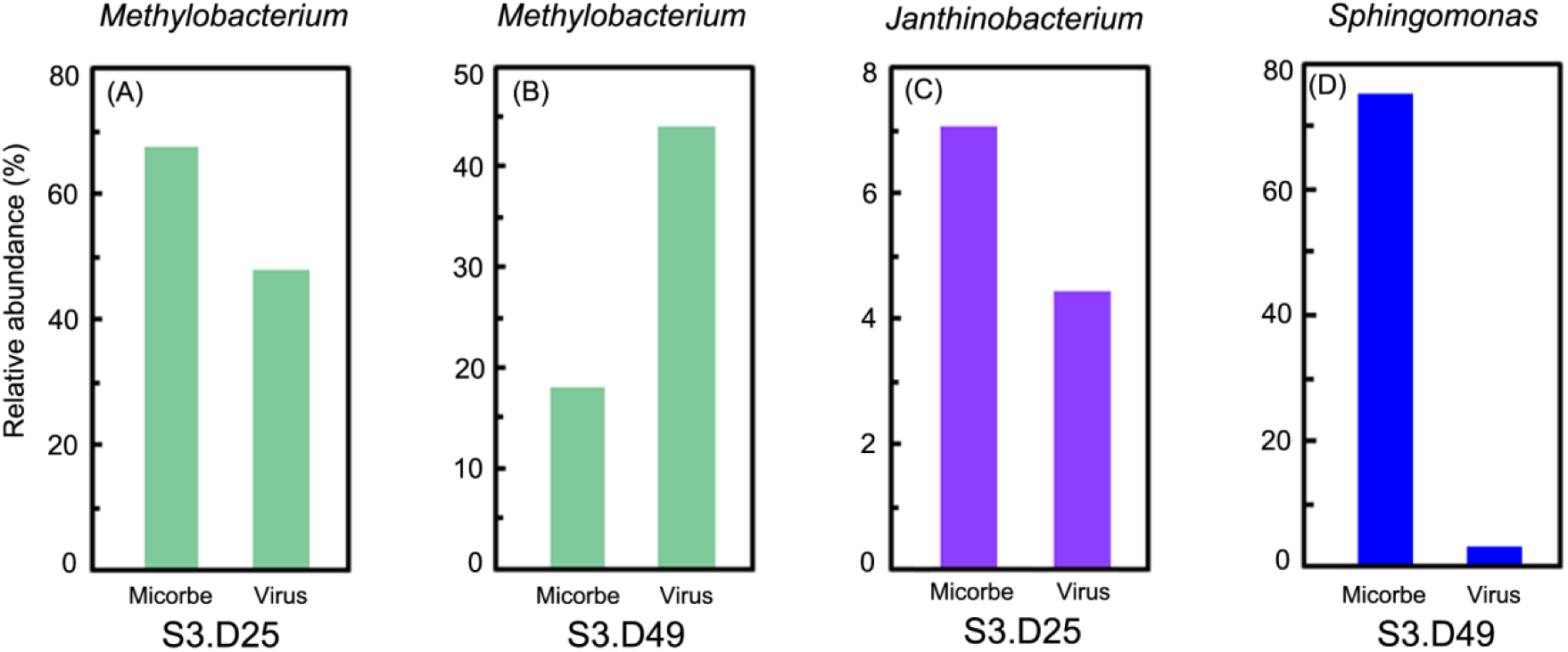
Relative abundances of three abundant (>1.0%) microbial genera and their viruses. (**A**) *Methylobacterium* in S3.D25; (**B**) *Methylobacterium* in S3.D49; (**C**) *Janthinobacterium* in S3.D25; (**D**) *Sphingomonas* in S3.D49. Relative abundances of microbes are based on 16S rRNA amplicon sequencing, and viral populations are based on their coverages generated by mapping quality-controlled reads to viral populations. Viruses were linked to hosts *in-silico* by three methods: Blastn, VirHostMatcher, and CRISPR matches.

## Conclusion

Glacier ice harbors diverse microbes, yet the associated viruses and their impacts on ice microbiomes have been unexplored. Glaciers around the world are rapidly shrinking, primarily due to the anthropogenic-enhanced warming of Earth’s ocean-atmosphere system (72), sand this will release glacial microbes and viruses that have been trapped and preserved for tens to hundreds of thousands of years (73, 74). At a minimum, this could lead to the loss of microbial and viral archives that could be diagnostic and informative of past Earth climate regimes; however, in a worst-case scenario, this ice melt could release pathogens into the environment. Starting with previously proposed robust *in silico* decontamination methods of glacier-ice microbial study (12), we further improved our procedures to remove microbial and viral contaminants on the ice core surfaces, and applied these methods to investigate microbial and viral communities archived in glacier ice cores. These data begin to fill the information gap on viruses archived in glacier ice and shed light on the potential impacts of viruses on their microbial hosts when active. Future studies will provide a better understanding of microbial and viral evolution and interactions, and will contribute to establishing predictive ecological models of past climate changes from such “frozen archive” environments.

## Materials and Methods

### Sterile artificial ice core sections and mock “contaminants”

An artificial ice core was constructed from sterile water, which was pre-filtered through a Millipore system (Cat No. MPGP04001, MillipakR Express 40 Filter, Merck KGaA) outfitted with a 0.22-μm mesh final filter and autoclaved at 121°C for 30 min, then frozen at −34°C for 12–24 hr in a 2L sterile plastic cylinder (Nalgene). The cylinder was transferred from −34°C to −5°C and kept at that temperature overnight to reduce the possibility of fracturing (which is caused by sudden temperature changes) before placing it at room temperature for about 30 min to melt the surface ice and expose the underlying ice core.

*Cellulophaga baltica* strain #18 (CBA 18; NCBI accession No. CP009976) was cultured in MLB medium [15 g sea salts (Cat No. S9883, Sigma), 0.5 g bacto peptone, 0.5 g yeast extract, 0.5 g casamino acids, 3 ml glycerol, and 1000 ml water] stationary overnight at room temperature. The cell concentration was measured by epifluorescence microscopy after the cells were captured on a 0.22-μm-pore-sized filter (Cat No. GTTP02500, Isopore) and stained by SYBR Green (Cat No. S9430, Sigma) as described previously (75) with some modifications. Briefly, cells on the filter were covered with several drops of 20×SYBR Green (Cat No. S11494, Life Technologies). After 15 min of staining in the dark, the SYBR Green was carefully removed with a 50-μl pipette and by touching the backside of the membrane with a Kimwipe (Kimtech). The filter was mounted on a glass slide with freshly-made anti-fade solution (1 mg ascorbic acid : 100 μl PBS : 100 μl glycerol) and a 25 mm^2^ cover slip. Cells on the filter were counted using epifluorescence microscopy (Zeiss Axio Imager.D2) with >350 cells or >20 fields counted, which was a reliable threshold to estimate the total bacterial abundance (76).

*Pseudoalteromonas* phages strain PSA-HP1 (NCBI: txid134839) were harvested from 95% lysed plaque assays (agar overlay technique). The concentration of PSA-HP1 was counted by a wet-mount method using SYBR Gold (Cat No. S11494, Life Technologies) staining and glass beads as described previously (52). The lambda phage DNA (100 μg/ml; 1.88×10^9^ copies/μl; genome size 4.8 kb) was purchased from Life Technologies (Cat. No. P7589). The above components (i.e., CBA 18, PSA-HP1, and lambda phage DNA) were combined in 1 ml ddH2O, which contained 1.00×10^6^ cells, 4.48×10^7^ viruses, and 1.88×10^8^ copies of lambda DNA to make the mock contaminants. The 1-ml mixtures were spread evenly on the artificial ice core surface with sterile gloved hands. The ice core was cut into three equal sized sections with a sterilized band saw, which was previously wiped with 75% ethanol and exposed to UV light for >12 hrs.

### Surface decontamination procedures

The decontamination procedure consisted of three steps (Fig. 1) following a previously published method (35) with slight modifications. First, the exterior (∼0.5 cm of the core radius) of the ice core was scraped away using a sterile band saw; second, the ice core was rinsed with 95% ethanol (v/v; Cat No. 04355223, Decon Labs) to remove another ∼0.5 cm of the ice core surface; third, a final ∼0.5 cm of the ice core surface was washed away with sterile water (Fig. 1; Fig. S1). After about 1.5 cm of the core surface was removed, the inner ice was the “clean” sample and collected for further analyses.

Two artificial ice core sections (Section 1 & 2) were processed using the decontamination procedure described above (Fig. 1B). The ice removed by the saw scraping (first step), water washing (third step), and the inner ice were collected as three different samples in sterile beakers. As a positive control, another ice core section was placed in a sterile beaker, which was not decontaminated (Fig. 1B). All sampling steps were conducted in a cold room (−5°C), which was exposed to UV light >12 hr before ice core processing to kill microbes and viruses in the air and on the surface of the instruments (e.g., band saw, washing systems, and flow hood; Fig. S1). In addition, we performed the washings with 95% ethanol and water in the BioGard laminar flow hood (Baker Company, model B6000-1) to avoid environmental contaminations (Fig. S1). Ice samples were melted at room temperature. One ml of each melted sample was preserved at 4°C and used for nested PCR to detect the coated lambda DNA (see Methods below). Other volumes of each sample were subjected to concentrating the microbes and viruses using 100 kDa Amicon Ultra Concentrators (EMD Millipore, Darmstadt, Germany). Each sample was concentrated to 0.8 ml and used for DNA extraction (see Methods below).

### Guliya ice core sampling and physiochemical conditions

The shallow plateau core (34.5 m depth) (∼PS; 35°14’ N; 81°28’ E; 6200 m asl) and the summit core 3 to bedrock (51.86 m depth) (S3; 35°17’ N; 81°29’ E, ∼6650 m asl) were drilled on the Guliya ice cap in 1992 and 2015, respectively (Fig. 3). Both cores were 10 cm in diameter and the bedrock temperature at the S3 site was about −15°C (77). Ice core sections (∼1 m each) were sealed in plastic tubing, placed in cardboard tubes covered with aluminum, and transferred at −20°C by truck from the drill sites to freezers in Lhasa, by airplane to freezers in Beijing, by airplane to Chicago and then by freezer truck to the Byrd Polar and Climate Research Center at the Ohio State University where they were stored at −34°C. The ice core sections were transferred from −34°C to the sampling room and kept at −5°C overnight to reduce the possibility of fracturing during surface decontamination by cutting and washing. Five samples were collected from the PS core at depths of 13.34–13.50 (sample name PS.D13.3), 13.50–13.67 (PS.D13.5), 24.12–24.54 (PS.D24.1), 33.37–33.52 (PS.D33.3), and 34.31–34.45 (PS.D34.3) meters (Fig. 3; Table S2). These ice samples were decontaminated using the surface-decontamination procedure described above, and the inner ice was collected for further analysis. In addition, the ice removed from the saw scraping and water washing was also collected for two samples (PS.D13.3 and PS.D13.5) as described for the artificial ice core sections, in order to evaluate the surface decontamination procedures using authentic ice samples. The microbial populations from two of the S3 core samples (S3.D41 and S3.D49) were published previously (12). Another sample S3.D25 (25.23–25.79 meters depth; not published) was collected at the same time as the two samples mentioned above and was included in this study (Fig. 3).

Four controls were used to trace possible sources of background contamination during ice sample processing as described previously (12). First, we assessed what microorganisms inhabited the air of the cold room laboratory in which the sampling took place. Cells from about 28 m^3^ of air were collected over four days of continuous sampling in the room using an air sampler (SKC Inc.) as described previously (12), during which the ice samples were processed at the same time. This provided an evaluation of the background contamination due to ice exposure to air during the processing (Sample AirColdRoom). Second, an artificial ice core was made from sterile water (same as above) frozen at − 34°C for 12–24 hr. This sterile core was processed in parallel with the authentic ice core samples through the entire analysis. This control allowed evaluation of contamination from the instruments used to process the ice (Sample ArtificialIce). Third, a blank control was established by extracting DNA directly from 300 ml of sterile water. This control allowed evaluation of contamination downstream of the ice processing, including the molecular procedures (DNA extraction, PCR, library preparation, and sequencing; Sample Blank). Finally, 30 μl of filtered and autoclaved water was subjected to standard 16S rRNA gene amplicon sequencing to check contamination from the sequencing procedures (Sample BlankSequencing).

A total of 300 ml of artificial ice, 300 ml of the blank control, and 100–300 ml each of above glacier ice samples were filtered through sterilized polycarbonate 0.22-μm-pore-sized filters (Cat No. GTTP02500, Isopore) to collect microbes including all bacterial/archaeal cells larger than 0.22 μm. The filters were preserved at −20°C until DNA extraction. Viruses in the filtrate of two samples (S3.D25 and S3.D49) were concentrated to 0.8 ml using 100 kDa Amicon Ultra Concentrators (EMD Millipore, Darmstadt, Germany) and preserved at 4°C until DNA extraction. To check for possible cross contamination among samples and potential viral contaminants introduced to the samples during processing, 1 ml of 0.22-μm-pore-size filtrate from the water of the Olentangy River (named RiverV; 39°59’52’’ N, 83°1’24’’ W, Columbus, Ohio) was co-processed in parallel with samples S3.D25 and S3.D49 throughout the entire analyses. All the biological work in this study after the ice sampling in the cold room laboratory was performed in an ultra-clean room, designed for experiments with low-biomass samples.

Dust, chemical ions, and oxygen isotopes of glacier ice were analyzed as described previously (78). The chronology of the ice cores, from which the samples were collected, was established using a combination of annual layer counting, ^14^C AMS dating of englacial plant fragments, and δ^18^Oatm age (L. G. Thompson, E. Mosley-Thompson et al., unpublished data).

### Genomic DNA extraction

The viral concentrates of samples S3.D25, S3.D49, and RiverV were subjected to isolating genomic DNA as previously described (79). Briefly, viral concentrates were treated with DNase I (100 U/ml) to eliminate free DNA, followed by the addition of 100 mM EDTA/100 mM EGTA to halt DNase activity; Genomic DNA was then extracted using Wizard® PCR Preps DNA Purification Resin and Minicolumns (Cat. No. A7181 and A7211, respectively; Promega, USA) (79, 80). Viral abundance was obtained by enumerating and comparing the counts of VLPs and beads (with a known concentration) using the wet-mount method (52).

Genomic DNA from all other samples was isolated with a DNeasy Blood & Tissue Kit (Cat No. 69506, QIAGEN) according to the manufacturer’s instructions, with an additional step of beating with beads to disrupt bacterial spores and Gram-positive cells before cell lysis by homogenizing at 3400 RPM for 1 min with 100 mg of autoclaved (121°C for 30 min) 0.1-mm-diameter glass beads (Cat No. 13118-400, QIAGEN) in a MiniBeadBeater-16 (Model 607, BioSpec Products).

### Nested PCR

Nested PCR experiments (81) were performed with two pairs of designed primers to detect lambda phage DNA in the artificial ice section samples, which were used to establish the clean surface decontamination procedures. The external primer set LamouterF (5’-CAACTACACGGCTCACCTGT-3’) and LamouterR (5’-ACGGAACGAGATTTCCGCTT-3’) amplifies a 674 bp fragment, and the nested primer set LaminnerF (5’-GAAGCTGCATGTGCTGGAAG-3’) and LaminnerR (5’-CACACTCTGGAGAGCACCAC-3’) amplifies a 189 bp fragment within the previous fragment. In the first PCR with the external primer sets, the 25 μl reaction mixture consisted of 12.5 μl 2× commercial mix (Cat No. M712B, GoTaq^®^ Green Master Mix, Promega), 1.25 μl of each external primer (LamouterF/LamouterR, 10 uM), 5.0 μl template DNA, and 5 μl of ddH2O. The amplification included a 5 min denaturation step at 95°C, followed by 40 cycles of 30 sec at 95°C, 30 sec at 56°C, and 50 sec at 72°C, with a final extension of 5 min at 72°C. For the nested PCR, the reaction mixture was identical to the first PCR, except that 5.0 μl of the first PCR product and 1.25 μl of each nested primer (LaminnerF/LaminnerR, 10 μΜ) were included. The amplification conditions were also identical to the first PCR except for the extension time of 20 sec at 72°C for 40 cycles of amplifications.

For each of artificial ice section samples (i.e., Cut1, Wash1, Inner1, Cut2, Wash2, Inner2, and Mix; Fig. 1B), 5 μl of melt water served as the DNA template in the first PCR. In addition, nested PCRs were performed using diluted lambda DNA (1.88×10^4^, 10^3^, 10^2^, 10^1^, 10^0^, and 10^−1^ copies, respectively) as templates to serve as a reference. A negative control was conducted with 5 μl of ddH2O as template.

### Real-time quantitative polymerase chain reaction (qPCR)

Each 20-μl reaction for qPCRs contained: 10 μl of 2× QuantiTect SYBR Green PCR Master Mix (Cat No. 204143, QIAGEN), 0.5 μl of each primer (10 μM), 3 μl of template DNA and 6 μl of RNase-free water. All reactions were performed in triplicate, using an Illumina Eco cycler (Cat No. 1010180).

Total bacterial and archaeal biomasses of the glacier ice samples and the “background” controls were estimated using qPCR after isolating DNA. The primer set 1406f (5′-GYACWCACCGCCCGT-3′) and 1525r (5′-AAGGAGGTGWTCCARCC-3′) was used to amplify bacterial and archaeal 16S rRNA genes (82). Thermocycling consisted of an initial polymerase activation and template DNA denaturation step at 95°C for 15 min, followed by 40 cycles of 95°C for 15 sec, 55°C for 30 sec, and 72°C for 15 sec. A standard curve was generated with a PCR product using primers 1406f/1525r from *Cellulophaga baltica* strain 18 (NCBI accession number of the complete genome, CP009976).

Total numbers of CBA 18 in each of artificial ice section samples (i.e., Cut1, Wash1, Inner1, Cut2, Wash2, Inner2, and Mix; Fig. 1B) were quantified using the primer set Cbal18M666_05390F (5′-ACGTACAAATAAGGAGAATGGCTT-3′) and Cbal18M666_05390R (5′-AGCGCTAATCCCTGTTGAGA-3′), which specifically targets a 61 bp fragment of an ATP synthase subunit C of CBA 18, with thermocycling: 95°C for 15 min, 45 cycles of 95°C for 15 sec, 60°C for 30 sec, and 70°C for 25 sec. Similarly, total PSA-HP1 numbers of these samples were quantified using strain-designed primer set 10-94a_dF (5′-TCTCTCGTCTTAATGACTTTCATCAT-3′) and 10-94a_dR (5′-TTCTTTCTCAACTTCCTGCTCTAA-3′), with the identical thermocycling conditions except that 50 cycles of amplifications were conducted. The standard curves of the above two qPCRs were generated with the PCR products from their primer sets and strains, respectively.

### Tag-encoded amplicon sequencing of the microbial community

Bar-coded primers 515f/806r (83) were used to amplify the V4 hypervariable regions of 16S rRNA genes of bacteria and archaea for all the glacier ice samples and the “background” controls. Resulting amplicons were sequenced by the Illumina MiSeq platform (paired-end reads), as described previously (83, 84). These experiments were performed at Argonne National Laboratory.

### Amplicon sequence analysis

Sequences with an expected error >1.0 or length <245 nt were excluded from the analyses (85). The remaining sequences were truncated to a constant length (245 nt). Various analyses were conducted using the QIIME (Quantitative Insights Into Microbial Ecology, version 1.9.1) software package (86) using default parameters, except that chimera filtering, operational taxonomic unit (OTU) clustering, and singleton excluding were performed with QIIME through the UPARSE pipeline (85). A phylogenetic tree was constructed with a set of sequence representatives of the OTUs using the method of FastTree (87). Chimeras were identified and filtered by UPARSE with the UCHIME algorithm using the ChimeraSlayer reference database (88), which is considered to be sensitive and quick (89). Reads were clustered into OTUs at 97% sequence similarity by UPARSE. A representative sequence from each OTU was selected for taxonomic annotation using the Ribosomal Database Project (RDP) classifier (90) from the RDP Release 11.5 database. Taxonomic assignments with <80% confidence were marked as unclassified taxa. Mitochondrial and chloroplast sequences were excluded from further analyses. A new profile of OTU composition for the ice samples was generated after *in silico* and proportional decontamination using R-OTU values >0.01 according to the method established previously (12). Briefly, an R-OTU value was designated as the ratio between the mean “absolute” abundance of OTUs in “background” controls and ice samples; then an approximated estimation of the “absolute” abundance of OTUs was calculated by multiplying the relative abundance of each OTU by the 16S rRNA gene copy number in a given sample (determined by qPCR). The OTUs with R-OTU values >0.01 were considered to be contaminants and were removed from the ice samples. Relative abundance of the microbial profiles was generated at genus and class levels. Principal coordinates analysis (PCoA) using weighted UniFrac metrics was performed to distinguish general distribution patterns of microbial profiles among all samples. The Mantel tests were conducted to evaluate the linkage between the microbial community structure and environmental parameters. The significance of the difference in microbial community between grouped samples (PS versus S3 core samples) was evaluated by Analysis of similarity statistics (ANOSIM) (91), which was performed using functions in the Vegan package version 2.4-4 (92) in R version 3.4.2 (93).

### Metagenomic sequencing of viral metagenomic DNA

The viral genomic DNA from three samples (S3.D25, S3.D49, and RiverV) was subjected to low-input library preparation pipeline using the Nextera^®^ XT Library Prep Kit (Cat No. 15032354, Illumina) in the ultra-clean room, according to our methods described previously (53–55). The metagenomes were sequenced by Illumina HiSeq 2000 platform (1×100 bp) at JP Sulzberger Genome Center at Columbia University.

### Viromic analysis and characterization of viral communities

All metagenomic analyses were supported by the Ohio Supercomputer Center (94). Virome sequence data were processed using iVirus pipeline with default parameters described previously (28, 51). Briefly, raw reads of three viromes, including two glacier ice samples (S3.D25 and S3.D49) and the River water control (RiverV), were filtered for quality using Trimmomatic v0.36 (95), followed by the assembly using metaSPAdes v3.11.1 (k-mer values include 21, 33, and 55) (96), and the prediction of viral contigs using VirSorter v1.0.3 in virome decontamination mode on Cyverse (56). The viral contigs (Categories 1, 2, 4, and 5) were first checked for contaminants by comparing them to some viral genomes considered as putative laboratory contaminants (e.g., phages cultivated in our lab including *Synechococcus* phages, *Cellulophaga* phages, and *Pseudoalteromonas* phages) using Blastn. Then they were clustered into populations if viral contigs shared ≥95% nucleotide identity across 80% of their lengths as described previously (28, 97). The longest contig within each population was selected as the seed sequence to represent that population. A coverage table of each viral population was generated using BowtieBatch and Read2RefMapper by mapping quality-controlled reads to viral populations and the resulting coverages were normalized by library size to per gigabase of virome (51). Rarefaction curves of the two glacier ice viromes were produced by changing viral population (length ≥10kb) numbers along sequencing depth (i.e., read number), which was obtained by subsampling quality-controlled reads (Fig. S3).

A total of 33 and 107 viral populations (length ≥10 kb) were obtained for two glacier ice samples (S3.D25 and S3.D49) and the river water control (RiverV) viromes, respectively. Mapping the quality-controlled reads of 3 viromes to 140 viral populations (33+107) found that the viral communities in the glacier ice samples were completely different from those in the river water control (Fig. S4), suggesting that the procedures for handling glacier ice samples were “clean” and no cross contamination was detected among these samples. Only the two glacier ice viromes were used for additional analyses.

Taxonomy assignments were performed using vConTACT v2.0 (60, 61). Briefly, this analysis compared the viral populations in this study to 2,304 viral genomes in the National Center for Biotechnology Information (NCBI) RefSeq database (release v85), and generated viral clusters (VCs) approximately equivalent to known viral genera (31, 60, 61, 63). The putative virus–host linkages were predicted *in silico* using three methods based on: i) nucleotide sequence composition, ii) nucleotide sequence similarity, and iii) CRISPR spacer matches, as described previously (31, 64). Thirty-three viral populations from glacier ice samples were linked to their microbial hosts using the oligonucleotide frequency dissimilarity (VirHostMatcher) measure, with ∼32,000 bacterial and archaeal genomes as the host database and a dissimilarity score ≤0.1 and possibility ≥80% as the threshold to pick the host (70). In addition to sequence composition analysis using VirHostMatcher, the nucleotide sequence of each viral population was compared (Blastn) to bacterial and archaeal genomes from the NCBI RefSeq database (release v81) and the database (∼32,000 genomes) used above. The viral sequences were considered for successful host predictions if they had a bit score of ≥50, E-value of ≤10^−3^, and average nucleotide identity of ≥70% across ≥2,000 bp with the host genomes (31). Finally, nucleotide sequences of 33 populations were compared to CRISPR spacers of bacterial and archaeal genomes in both databases using the sequence similarity method. The CRISPR spacers were identified using MinCED (mining CRISPRs in environmental data sets; http://github.com/ctSkennerton/minced), which is a derivative of CRT (CRISPR Recognition Tool) (98), and compared to nucleotide sequences of 33 viral populations. Hosts were selected if the spacers had zero mismatches to viral populations.

## Data availability

The amplicon sequences obtained in this study have been deposited in the NCBI Sequence Read Archive under BioProject accession number PRJNA594142. The viral metagenomes are available through iVirus (https://datacommons.cyverse.org/browse/iplant/home/shared/iVirus/Tibet_Glacier_viromes_2017), including raw and quality-controlled reads and viral populations.

## Supplemental material

Supplemental material for this article is available at https://doi.org/10.6084/m9.figshare.11427246.v1

## Acknowledgements

The 2015 Guliya ice cores were collected and analyzed as part of a collaborative expedition between The Ohio State University’s Byrd Polar and Climate Research Center (BPCRC) and the Institute of Tibetan Plateau Research of the Chinese Academy of Sciences, funded by the National Science Foundation’s Paleoclimate Program award #1502919 and the Chinese Academy of Sciences, respectively. Partial support for Dr. Zhong was provided by a Gordon and Betty Moore Foundation Investigator Award to MBS (#3790), by NSF’s Paleoclimate Program award #1502919, and the BPCRC Postdoctoral Program. This is BPCRC contribution number 1586. The work conducted by the U.S. Department of Energy Joint Genome Institute is supported by the Office of Science of the U.S. Department of Energy under contract no. DE-AC02-05CH11231. The authors greatly appreciate the help provided by Benjamin Bolduc, Ho Bin Jang, Ann C. Gregory, Gareth Trubl, and Dean R. Vik with virome analysis, by Donald V. Kenny and Ping-Nan Lin with ice core sampling, by Paul Green, Emilie Beaudon, and M. Roxana Sierra-Hernández with construction of the surface decontamination system, and by the Sullivan, Thompsons, and Rich laboratories for critical review and comments through the years. Bioinformatics were supported by the Ohio Supercomputer Center.

